# IgG3 subclass antibodies recognize antigenically drifted influenza viruses and SARS-CoV-2 variants through efficient bivalent binding

**DOI:** 10.1101/2022.09.27.509738

**Authors:** Marcus J. Bolton, Claudia P. Arevalo, Trevor Griesman, Shuk Hang Li, Paul Bates, Patrick C. Wilson, Scott E. Hensley

## Abstract

The constant domains of antibodies are important for effector functions, but less is known about how they can affect binding and neutralization of viruses. Here we evaluated a panel of human influenza virus monoclonal antibodies (mAbs) expressed as IgG1, IgG2 or IgG3. We found that many influenza virus-specific mAbs have altered binding and neutralization capacity depending on the IgG subclass encoded, and that these differences result from unique bivalency capacities of the subclasses. Importantly, subclass differences in antibody binding and neutralization were greatest when the affinity for the target antigen was reduced through antigenic mismatch. We found that antibodies expressed as IgG3 bound and neutralized antigenically drifted influenza viruses more effectively. We obtained similar results using a panel of SARS-CoV-2-specific mAbs and the antigenically advanced B.1.351 strain of SARS-CoV-2. We found that a licensed therapeutic mAb retained neutralization breadth against SARS-CoV-2 variants when expressed as IgG3, but not IgG1. These data highlight that IgG subclasses are not only important for fine-tuning effector functionality, but also for binding and neutralization of antigenically drifted viruses.

**Significance:** Influenza viruses and coronaviruses undergo continuous change, successfully evading human antibodies elicited from prior infections or vaccinations. It is important to identify features that allow antibodies to bind with increased breadth. Here we examined the effect that different IgG subclasses have on monoclonal antibody binding and neutralization. We show that IgG subclass is a determinant of antibody breadth, with IgG3 affording increased neutralization of antigenically drifted variants of influenza virus and SARS-CoV-2. Future studies should evaluate IgG3 therapeutic antibodies and vaccination strategies or adjuvants that may skew antibody responses toward broadly reactive isotypes.

---

Antibodies are key components of effective immune responses against viruses. Antibody responses can be fine-tuned in germinal centers, with changes in specificity or affinity being mediated by somatic hypermutation in the antibody variable domains of B cells (1, 2). Isotype or subclass switching, which swaps in a different cassette of heavy chain constant domains, can also occur in germinal center B cells, and this can alter antibody effector functions (reviewed in (3)). Different antibody isotypes vary in degree of complement binding, antibody-dependent cellular cytotoxicity (ADCC) or phagocytosis (ADCP).

It is commonly thought that isotype switching does not affect antigen binding, since the constant domains of antibodies are distal to the variable antigen binding domains. However, immunoglobulin heavy chain constant domains can have allosteric effects on antigen recognition (reviewed in (4, 5)). For example, studies have shown that an anti-tubulin antibody expressed as IgG1 and IgA1 had differences in binding affinity (6), and that the fine specificity of a *Cryptococcus neoformans-reactive* antibody changed when the murine IgG1 isotype was swapped for human IgM or IgG3 (7). Interestingly, isotype swapping of some monoclonal antibodies (mAbs) can affect affinity and specificity, but this is not an ubiquitous phenomenon for all mAbs (5). Less is known about how isotype variation affects antigen recognition of antiviral antibodies. Most human antiviral mAb studies include sequencing of the antibody variable domains, but not antibody constant domains, from plasmablasts following infection and vaccination. Antibody variable domains are then typically cloned into IgG1 vectors and mAbs are expressed as IgG1 antibodies (8). This streamlined process allows for the study of many unique antibody clones, but does not account for the naturally encoded isotype of each mAb. More recent technologies allow for rapid sequencing of both the antibody variable and constant domains of antiviral antibodies (9).

Viruses that rapidly change over time, such as influenza viruses, HIV-1, and coronaviruses, have the ability to escape previously generated antibody responses (reviewed in (10–12)). Influenza viruses continuously acquire substitutions in external hemagglutinin (HA) and neuraminidase proteins which necessitates continual reformulation of annual vaccines. Coronaviruses, like 229E, have also been shown to undergo significant antigenic change over time (13). The role of IgG subclass in the reactivity to antigenically distinct strains has been studied in the context of HIV-1-specific polyclonal serum (14). These studies demonstrated that serum fractions containing the individual subclasses had an ordered neutralization breadth of IgG3>IgG1>IgG2 to various HIV-1 strains (14). It is unclear if these findings are more widely applicable to other antiviral antibodies, or if these are unique findings confined to certain mAbs.

Here we completed a series of experiments to determine if the IgG subclass of antibodies affects antigen binding and virus neutralization of antigenically drifted viruses. First, we evaluated a panel of influenza-specific mAbs expressed as IgG1, IgG2, and IgG3. We then assessed the role of antibody valency by generating F(ab’)_2_ and F(ab) fragments of each mAb and we tested binding to matched and antigenically drifted influenza variants. Finally, we completed experiments using a panel of SARS-CoV-2 mAbs to determine if our influenza virus findings were broadly applicable to other virus types.

## Results

### IgG subclass confers subtle differences in HA binding and neutralization of influenza viruses when tested against antigenically matched viral strains

To assess the role of IgG subclass in the binding and neutralization of influenza viruses, we recombinantly expressed seven influenza virus HA-specific human mAbs with heavy chain constant domains of either IgG1, IgG2, or IgG3 (**Fig. 1A, Table S1**). We excluded IgG4 in this study due to its propensity to become functionally monovalent because of interchain instability (15), which can convolute comparative analyses with the other IgG subclasses. We chose mAbs that bind to different epitopes on the head and stalk domains of H1 and H3 HA proteins (**Fig. 1B**). We tested 5 mAbs that bind to epitopes in the HA head domain, including H3 antigenic site B (mAbs 10117-1B02, 10117-3A06, and 10040-4E01), H3 antigenic site E (mAb 10053-1G05), and H1 antigenic site Sa (mAb EM-4C04) (16–19). We tested 2 mAbs, 70-1F02 and CR9114, that bind to the HA stalk domain. 70-1F02 binds to group 1 HAs and CR9114 has broad reactivity for group 1, group 2, and influenza B HAs (17, 20, 21). We observed subtle, but reproducible differences in HA ELISA binding for some of the IgG subclass swapped mAbs (**Fig. 1C**). Three of the seven mAbs (EM-4C04, 10053-1G05, and 10117-1B02) had reduced binding capacity when expressed as IgG2 (2–4-fold) and one mAb (10117-1B02) bound less efficiently when expressed as IgG3. We measured virus neutralization capacities of these same mAbs and differences among IgG subclasses mostly mirrored the ELISA binding results (**Fig. 1D**). These data indicate that IgG constant domains alone can confer minor differences in antigen binding and neutralization of influenza viruses for the different IgG subclasses.

**Fig. 1.**
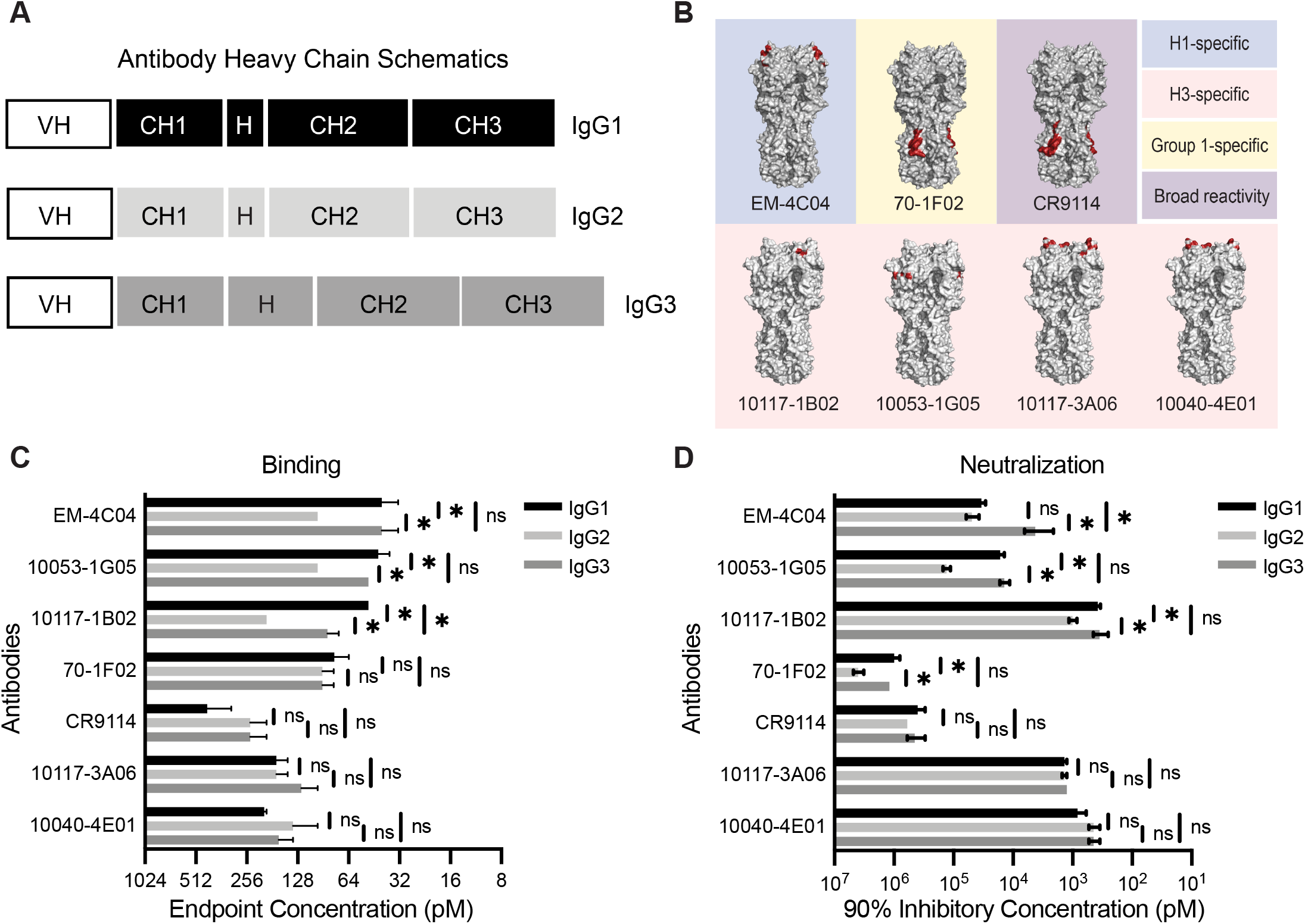
Influenza virus-specific antibodies display subtle changes in binding and neutralization upon IgG subclass swap. (**A**) Schematic of heavy chain domains of subclass-swapped mAbs. (**B**) The binding sites of mAbs are outlined red on HA crystal structures from representative H1 (PDB 3UBN) and H3 (PDB 4O5I) HAs visualized with PyMOL. (**C**) ELISAs were completed to measure binding of anti-influenza virus mAbs expressed as IgG1, IgG2, and IgG3. Endpoint titers were plotted as the mean, with error bars depicting the SEM of 3-4 independent experiments. Subclasses for each mAb were compared by one-way ANOVA with Tukey’s multiple comparisons post-test (*p <0.05). (**D**) Neutralization assays were completed for subclass-swapped mAbs. 90% inhibitory concentration values were plotted as the mean, with error bars depicting the SEM of 3-4 independent experiments. Subclasses for each mAb were compared by one-way ANOVA with Tukey’s multiple comparisons post-test. *p <0.05.

### Bivalent binding is needed for IgG subclass-dependent differences in binding and neutralization

We completed experiments to identify which of the antibody structural domains conferred subtle subclass-specific differences in binding and neutralization among our panel of influenza virus mAbs. We generated both F(ab’)_2_ and F(ab) fragments for three of the mAbs (10053-1G05, EM-4C04, and 10117-1B02) that displayed subtle subclass differences (**Fig. 2**, **Fig. S1**). F(ab’)_2_ fragments, which lack the 2^nd^ and 3^rd^ constant domains of the antibody heavy chain (which together comprise the Fc domain), were generated by cleavage of full-length antibody with the site-specific IgG protease, IdeS (**Fig. S1**). Like full-length antibodies, F(ab’)_2_ fragments are functionally bivalent because the hinge is left intact in this molecule. Unlike full-length antibodies and F(ab’)_2_ fragments, F(ab) fragments are monovalent, with the heavy chain being only comprised of the variable domain and 1^st^ constant domain (**Fig. S1**). We found that F(ab’)_2_ fragments for each of the subclasses bound antigen and neutralized virus to a similar degree as full-length antibody (**Fig. 2**). Conversely, F(ab) fragments had reduced binding and neutralization relative to full-length and F(ab’)_2_ fragments and we did not observe subclass-specific differences with F(ab) fragments (**Fig. 2**). This suggests that the antibody Fc domain does not contribute to subclass-specific differences in binding and neutralization. Unlike other antibody isotypes such as IgM and IgA which readily form multimers that increase valency, IgG is maintained as monomers, which have a fixed maximum valency of two (22, 23). Therefore, the differences that we observe with our subclass-swapped mAbs are likely due to bivalent binding capacity differences among the IgG subclasses mediated by differences in the hinge domain. These results are consistent with the relative differences in hinge flexibility between the IgG subclasses, where IgG3 > IgG1 > IgG2 (24, 25).

**Fig. 2.**
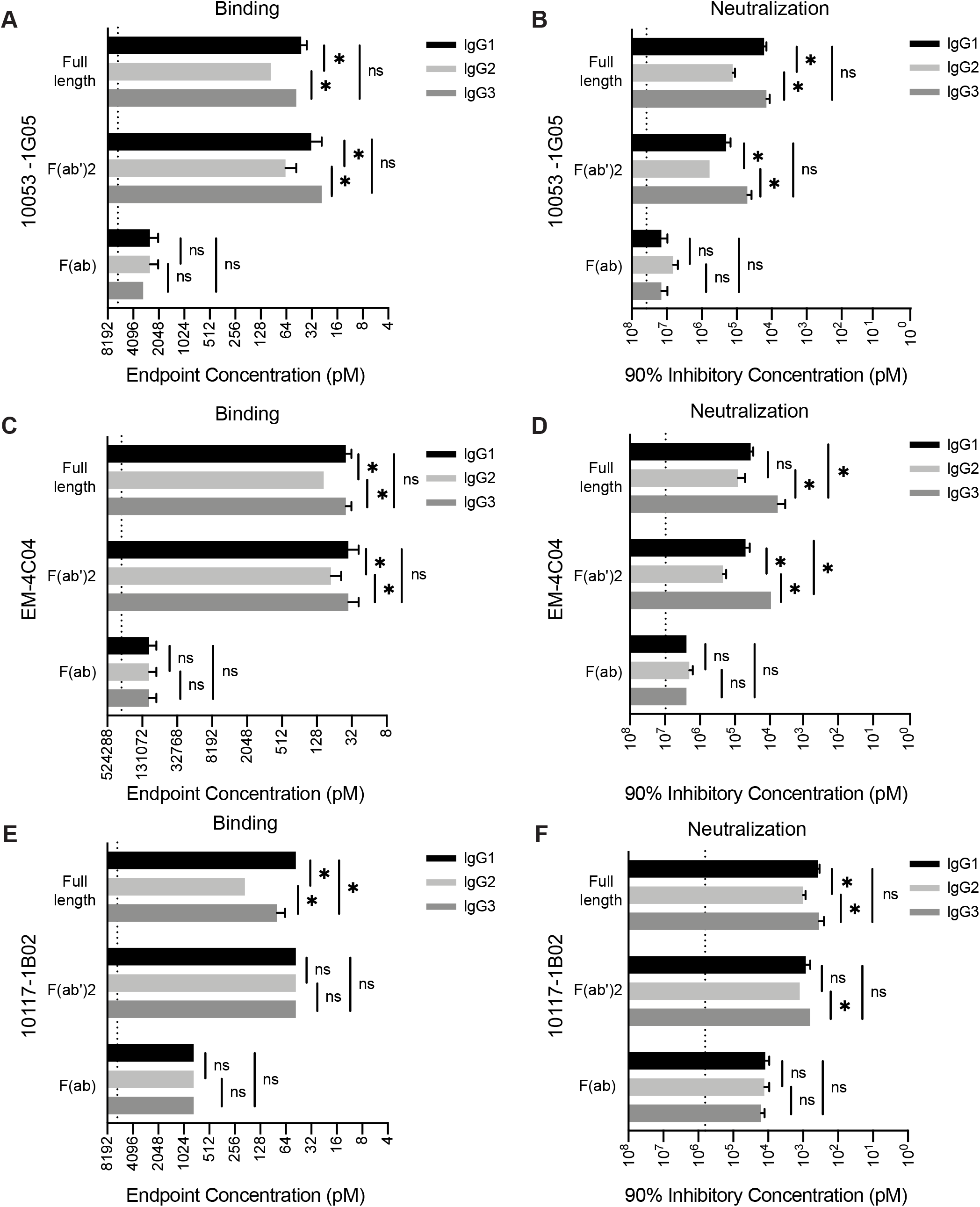
IgG subclass differences in binding and neutralization require bivalent binding. We completed experiments with IgG subclasses expressed as full-length, F(ab’)_2_, and F(ab) species. ELISA binding was measured for different antibody species of mAbs 10053-1G05 (**A**), EM-4C04 (**C**), and 10117-1B02 (**E**) expressed as IgG1, IgG2, and IgG3. Endpoint concentrations were plotted as the mean, and error bars indicate the SEM of 3-4 independent experiments. Statistical comparison of log2 transformed values for each mAb species were completed using a one-way ANOVA with Tukey’s multiple comparisons post-test. *p <0.05. Neutralization assays were completed for different antibody species of mAbs 10053-1G05 (**B**), EM-4C04 (**D**), and 10117-1B02 (**F**) expressed as IgG1, IgG2, and IgG3. Mean values of 90% inhibitory concentrations were plotted and error bars represent the SEM of 3-4 independent experiments. Statistical comparison of log2 transformed values for each mAb species were completed using a one-way ANOVA with Tukey’s multiple comparisons post-test. *p <0.05.

### IgG3 mAbs bind efficiently to antigenically drifted influenza virus antigens

Our data suggest that the differences in neutralization capacity we observe between the IgG subclasses depend on bivalent binding of the mAb, since monovalent F(ab)s yielded no differences among the subclasses. Due to an error-prone polymerase, influenza viruses are constantly evolving, often acquiring mutations that abrogate the binding of antibodies elicited by previous vaccinations or infections. When these changes occur in antibody epitopes, bivalent binding is often required for low affinity antibody recognition of antigenically drifted influenza antigens (18, 26–28). Therefore, we hypothesized that the small differences in binding and neutralization we observed among the subclasses would be exacerbated when measured against a target antigen that reduces the binding affinity of the mAb. To test this, we completed additional experiments with the 10053-1G05 mAb, which we previously found binds to antigenic site E of H3 (18). Antigenic site E is an important target of neutralizing antibodies and has undergone antigenic change in recent years (29). We tested binding and neutralization of IgG subclass-swapped versions of the 10053-1G05 mAb to two antigenically distinct viruses: A/Victoria/210/2009 (H3N2/2009) and A/Singapore/INFIMH-16-0019/2016 (H3N2/2016). Among the residues that differ between these two strains is a N121K substitution in HA site E (**Fig. 3A**). Consistent with our earlier experiments, all of the 10053-1G05 mAb subclasses efficiently bound and neutralized the H3N2/2009 virus, with a subtle reduction in binding and neutralization of the IgG2 version of the mAb (**Fig. 3B-C**). However, we observed large differences in the ability of the 10053-1G05 mAb subclasses to recognize and neutralize the antigenically drifted H3N2/2016 virus (**Fig. 3B-C**). The 10053-1G05 mAb expressed as IgG1 and IgG2 bound poorly to H3N2/2016 (endpoint concentrations ≥ 12.5 nM) and failed to neutralize this virus (IC_90_ ≥ 1200 nM), whereas the IgG3 version of the mAb efficiently recognized (endpoint concentration: 0.23 nM) and neutralized (IC_90_: 66.8 nM neutralization) H3N2/2016.

**Fig. 3.**
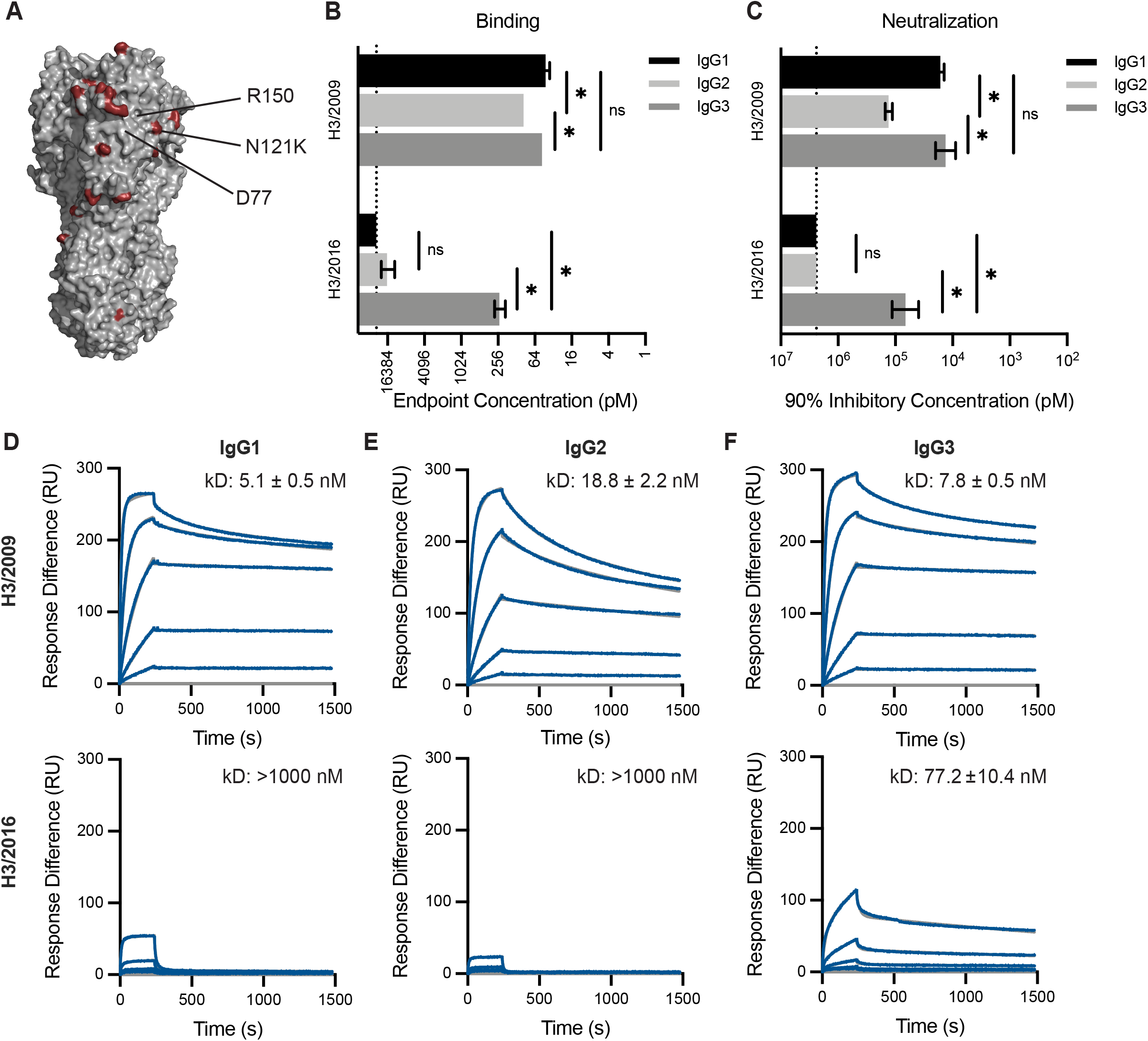
mAb 10053-1G05 retains binding and neutralization capacity to a drifted influenza virus strain when expressed as IgG3. (**A**) HA model (PDB 4O5I) depicting amino acid substitutions (red) that distinguish the 2009 and 2016 H3s. Known epitope residues of mAb 10053-1G05 are indicated. ELISAs (**B**) and neutralization assays (**C**) were completed with mAb 10053-1G05 expressed as different IgG subclasses. ELISA binding values are shown as the mean endpoint concentration and neutralization values are shown as the mean 90% inhibitory concentration, with error bars indicating the SEM for three independent experiments. Dotted line indicates the limit of detection for the assay. Statistical comparison of the IgG subclasses for each antigen/virus were completed using a one-way ANOVA on log2 transformed values with Tukey’s multiple comparisons post-test. *p<0.05 (**D-F**) SPR binding curves of 10053-1G05 IgG1, IgG2, and IgG3 to the 2009 H3 HA (top) and the 2016 H3 HA (bottom). Dissociation constants (kD) were determined from Langmuir (1:1 binding) model fits and are displayed as the mean ± SEM for 2-3 independent experiments.

To further confirm our findings, we used surface plasmon resonance (SPR) to measure the binding kinetics of mAb 10053-1G05 to HA proteins from both H3N2/2009 and H3N2/2016. Similar to our ELISA results, we found that 10053-1G05 expressed as IgG1 and IgG3 had similar binding affinities to H3N2/2009, with an average affinity (kD) of 5.1 nM and 7.8 nM, respectively (**Fig. 3D,F; top**). IgG2 had a lower affinity with an average kD of 18.8 nM when binding to H3N2/2009 (**Fig. 3E; top**). When we measured binding kinetics of 10053-1G05 IgG subclasses to the drifted HA, H3N2/2016, we found that only IgG3 bound appreciably (kD: 77.2 nM) (**Fig. 3F; bottom**), whereas the IgG1 and IgG2 forms of 10053-1G05 had no detectable binding (kD: <1000 nM) (**Fig. 3D,E; bottom**). In order to determine if these findings were generalizable to the other mAbs in our panel, we tested HA binding of additional mAbs against influenza drift and shift variants. We identified three more mAbs (10117-1B02, 10117-3A06, and CR9114), that bound to antigenically drifted/shifted variants most effectively when expressed as IgG3 (**Fig. S2**).

### SARS-CoV-2 mAbs bind and neutralize wild-type and variant strains more effectively as IgG3

Given our results that influenza virus-specific mAbs expressed as IgG3 bind and neutralize influenza drift variants more effectively, we completed additional experiments to determine if this was more widely applicable to other viruses that undergo antigenic change. SARS-CoV-2 rapidly diversified following its first detection in humans in 2019. While many distinct lineages emerged, we initially focused on the B.1.351 lineage of viruses, which possesses substitutions in the spike protein that disrupt the binding of monoclonal antibodies and polyclonal serum antibodies (30).

We produced SARS-CoV-2 mAbs with the same variable domains with either IgG1 or IgG3 constant domains. We focused on a previously characterized panel of neutralizing mAbs isolated from SARS-CoV-2 infected patients that bound to the SARS-CoV-2 receptor binding domain (RBD) of the spike protein (9). We characterized five mAbs that had variable reductions in binding to the RBD from B.1.351 lineage viruses (B.1.351 RBD) compared to that of an isolate from early in the pandemic (WA1 RBD) (**Fig. 4A**). Consistent with our studies of influenza virus mAbs, we found that the IgG3 form of four out of five SARS-CoV-2 mAbs bound to WA1 RBD more effectively compared to the IgG1 form of the same antibodies (**Fig. 4B**). All of the SARS-CoV-2 mAbs tested bound to the antigenically advanced B.1.351 RBD more effectively when expressed as IgG3 (**Fig. 4B**). The S144-1079 mAb is particularly interesting since the IgG1 and IgG3 forms bound similarly to the WA1 RBD but only the IgG3 form bound appreciably to the B.1.351 RBD (**Fig. 4B**). To measure the neutralization capacities of these subclass-swapped mAbs, we utilized a VSV-pseudotype neutralization assay. Consistent with the ELISA binding results, we found that many of the mAbs neutralized VSV expressing the SARS-CoV-2 spike more effectively when expressed as IgG3, with relative differences exceeding 100-fold with mAbs S24-223 and S20-74 against both viruses, and mAb S144-1079 against the virus expressing a B.1.351 spike (**Fig. 4C,D**). The S144-1079 mAb neutralized an early isolate of SARS-CoV-2 (WA1 strain) equally well when expressed as IgG1 or IgG3, but only neutralized virus with a spike of the antigenically advanced B.1.351 strain when expressed as IgG3 (**Fig. 4B-D**). We went on to test whether a licensed SARS-CoV-2 therapeutic mAb, REGN10933 (casirivimab) would yield similar results when expressed as IgG3 (**Fig. 4E-G**). We found that both IgG1 and IgG3 forms of REGN10933 efficiently neutralized WA1 pseudovirus (**Fig. 4E**), but REGEN10933 IgG3 was ≥100-fold more potent than REGEN10933 IgG1for neutralization of variant strains Beta (**Fig. 4F**) and Omicron (**Fig. 4G**). Together, these data mirror our findings for influenza mAbs where IgG3 mAbs bound and neutralized antigenically drifted viruses more efficiently than IgG1. This suggests that our findings are generally applicable to other antiviral mAbs that require bivalent binding.

**Fig. 4.**
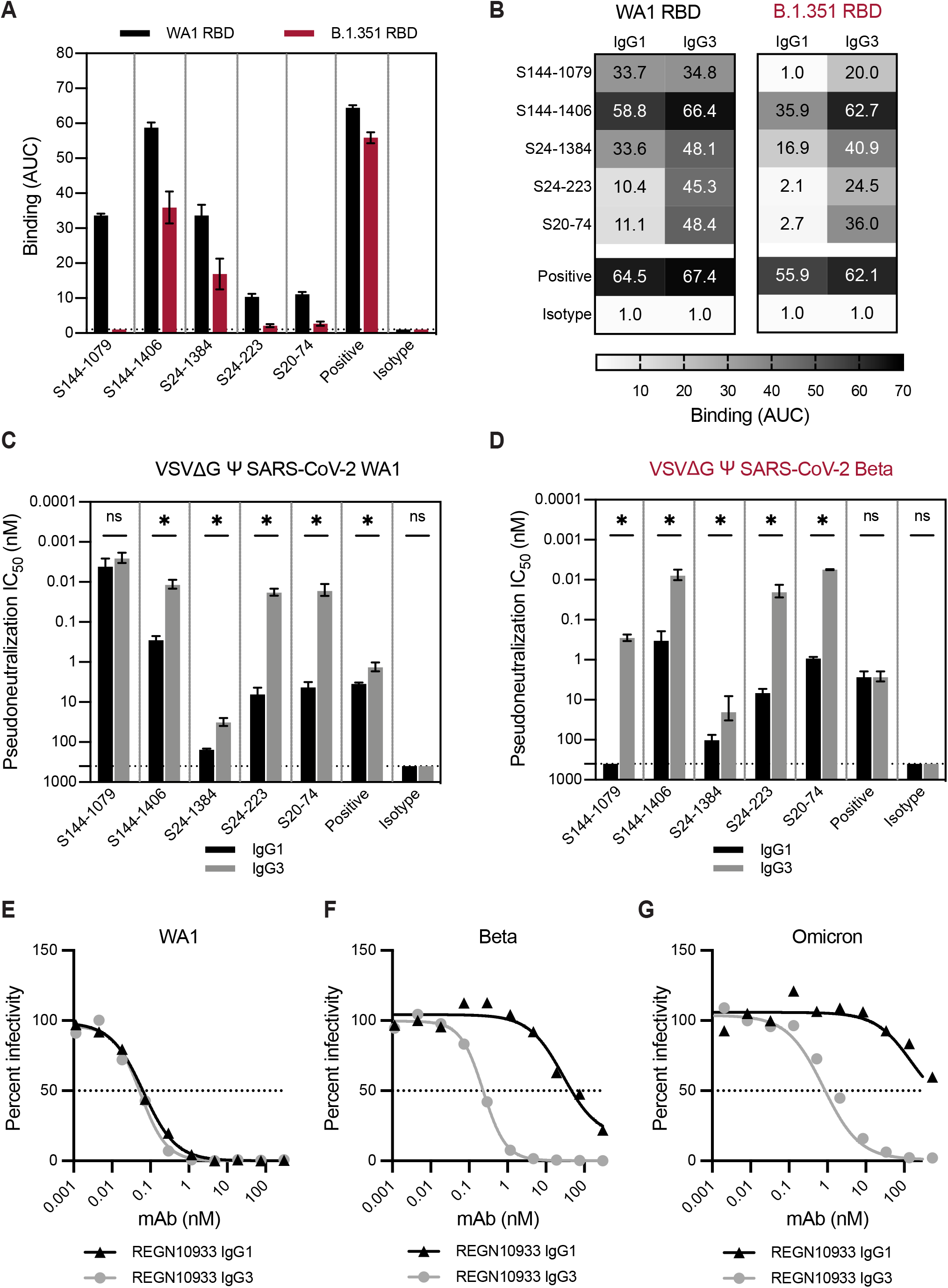
SARS-CoV-2 mAbs have greater binding and neutralization capacities when expressed as IgG3. A panel of SARS-CoV-2 mAbs were tested by ELISA (**A-B**) and *in vitro* neutralization assays (**C-D**). mAb S144-466 served as a positive control, and influenza mAb EM-4C04 served as an isotype control. (**A**) ELISAs were first completed with each mAb expressed as IgG1 against SARS-CoV-2 RBD proteins, WA1 (black) and B.1.351 (red). Dotted line indicates the limit of detection for the assay. Binding titers (AUC) are shown as the mean and error bars represent the SEM of three independent experiments. (**B**) ELISAs were then completed with each mAb expressed as IgG1 and IgG3 and the WA1 and B.1.351 RBDs. Binding titers (AUC) are shown as the mean of three independent experiments. (**C,D**) Neutralization assays were completed using mAbs expressed as IgG1 and IgG3 and VSV pseudotype viruses bearing SARS-CoV-2 WA1 spike (**C**) or the Beta (B.1.351) spike (**D**). Neutralization titers are shown as the mean, with error bars representing the SEM of 50% inhibitory concentrations for three independent experiments. (**E-G**) Neutralization assays were completed with therapeutic mAb REGN10933 expressed as IgG1 or IgG3 and VSV pseudotype viruses bearing SARS-CoV-2 spikes from WA1 (**E**), Beta (B.1.351) (F), or Omicron (BA.1) strains. Percent infectivity of wells (100 corresponding to virus-only control wells) was plotted against mAb concentration, and the dotted line at 50% is drawn to visualize IC_50_ values for mAbs. (**C,D**) Statistical comparisons between IgG1 and IgG3 for each mAb were completed using an unpaired t-test of log2 transformed values. *p<0.05

## DISCUSSION

In this study, we show that IgG constant domains affect the binding and neutralization capacity of both influenza virus and SARS-CoV-2 mAbs. When affinity for antigen is high, these differences are minor, but as the affinity for antigen is reduced through antigenic variation, we find that antibody constant domains can significantly alter binding and neutralization capacity. We found that these differences required bivalent antibody molecules, indicating the importance of antibody valency in mediating IgG subclass-specific differences. In particular, we found that antibodies expressed as IgG3, which have a long flexible hinge (31), usually bind better and neutralize antigenically drifted influenza viruses and SARS-CoV-2 viruses more efficiently compared to IgG1. These data are consistent with two recent studies that found enhanced neutralization potency for a broadly-reactive SARS-CoV-2 mAb when expressed as IgG3 (32, 33), and another that found that enhanced neutralization of an HIV-1 IgG3 mAb was dependent on the extended hinge length of IgG3 (31). Interestingly, the overall relative differences between IgG1 and IgG3 binding and neutralization in our studies were greater for the SARS-CoV-2 mAbs compared to influenza virus mAbs. It is possible that subclass differences play a larger role in the recognition of SARS-CoV-2 viruses compared to influenza viruses due to factors such as differences in virion glycoprotein density.

The increased cross-reactivity afforded by IgG3 could be a particularly useful feature for vaccination strategies aimed at providing protection against rapidly evolving pathogens. Our data suggest that antibody repertoires with large proportions of IgG3 might provide better protection against antigenically drifted viral strains. Subclass distribution following influenza virus or SARS-CoV-2 infections and vaccinations are typically dominated by IgG1 (34–38); however, it may be possible to develop new vaccination strategies and adjuvants to skew IgG subclass responses toward IgG3. Despite its apparent utility demonstrated *in vitro*, there are potential limitations to IgG3 as an effective molecule *in vivo*. IgG3 has a reduced half-life compared to the other IgG subclasses, due to its decreased affinity for neonatal Fc receptor and increased susceptibility to proteolytic cleavage (reviewed in (39)). Because of these reasons, our findings may prove to be more applicable to the therapeutic antibody field, where antibody engineering can overcome the natural limitations of IgG3, while retaining the attributes contributing to its broad reactivity reported here. Further studies should evaluate the *in vivo* therapeutic potential of IgG3 antibodies engineered to have longer half-lives.

Influenza viruses and coronaviruses will continue to present major public health burdens, highlighting the need for effective vaccines and therapeutics that elicit broad protection against virus variants. It is imperative that we understand the full scope of an effective antiviral antibody response, with breadth of binding and level of effector engagement being key factors. By further elucidating all of the functional changes antibodies acquire via the isotype or subclass they encode, we will continue to gain a more comprehensive image of antibody diversity within immune repertoires.

### Materials and Methods

#### Cell lines

293T cells were cultured in Dulbecco’s Minimal Essential Medium (DMEM) supplemented with 10% fetal bovine serum (FBS). MDCK-Siat1-TMPRSS2 cells were provided by Jesse Bloom (Fred Hutchinson Cancer Research Center) and cultured in Minimal Essential Medium (MEM) supplemented with 10% FBS. VeroE6/TMPRSS2 cells were provided by Stefan Pohlmann (German Primate Center, Leibniz Institute for Primate Research) and cultured in DMEM + 10% FBS (40). Adherent cell lines were maintained at 37°C in 5% CO_2_. Suspension 293F cells were maintained in Freestyle 293F Expression Medium at 37°C in 8% CO_2_.

## METHOD DETAILS

### Antibody heavy chain construct design

Expression plasmids encoding the human antibody IgG1 (IGHG1*03) heavy chain gene for each monoclonal (mAb) were obtained from sources indicated (**Supplementary Table 1**). IgG2 and IgG3 heavy chain constructs for each mAb were generated as previously described (41). Briefly, gBlock gene fragments (IDT, Coralville, IA, USA) encoding human IgG2 (IGHG2*01) or IgG3 (IGHG3*05) constant domains (CH1, hinge, CH2, and CH3) were cloned by Sal I and Hind III restriction digest into an AbVec-IgG1 mammalian expression vector (42). Monoclonal F(ab) expression plasmids for IgG1, IgG2, and IgG3 heavy chains were generated in a similar fashion to the full-length heavy chain constructs. For IgG1, a gene fragment encoding CH1 and terminating after the first cysteine in the hinge domain (..EPKSC*) was cloned into mAb expression vectors. For IgG2 and IgG3, gene fragments encoding only the CH1 domain were cloned into mAb expression vectors.

### Monoclonal antibody generation and purification

For monoclonal antibody transfections, 11 μg each of heavy and light chain plasmids were cotransfected in T-175 flasks of 293T cells using polyethylenimine in Opti-MEM. One day post-transfection, media was replaced with DMEM/F-12 supplemented with Nutridoma-SP. Cell culture supernatants were collected four days post-transfection and antibody was purified by affinity chromatography. Full-length antibodies were purified with protein A/G agarose, and F(ab)s were purified using CaptureSelect IgG-CH1 Affinity Matrix. All antibody species were eluted from affinity capture with the addition of IgG elution buffer, followed by neutralization with 1:10 eluent volume of 1.0 M Tris (pH 8.8). F(ab’)_2_ antibodies were produced by cleaving full-length antibody with the FragIT kit (bacterial protease, IdeS) according to manufacturer’s instruction. Antibodies were concentrated and buffer exchanged to PBS with Amicon Ultra centrifugal filters. Antibody concentrations were determined by measuring absorbance at 280 nm with a NanoDrop spectrophotometer, and molar concentrations were calculated for assays. The sequences of all plasmids encoding heavy and light chain domains were confirmed by Sanger sequencing prior to transfection. Antibody preparations were analyzed by reducing SDS-PAGE to check for protein integrity and absence of contaminants.

### Viruses

Influenza viruses A/Victoria/210/2009 H3N2 (X-187, GenBank HQ378745), A/California/07/2009 H1N1 (GenBank NC_026433.1) with a D225G substitution, and A/Singapore/INFIMH-16-0019/2016 H3N2 (GenBank MW298182.1) were generated by reverse genetics as previously described (43). All viruses were launched with cognate HA and NA genes, and internal segments from A/Puerto Rico/8/1934 H1N1. Each of the eight influenza virus segments in a pHW2000 vector were co-transfected using Lipofectamine 2000 in a co-culture of MDCK and 293T cells. The following day, media was replaced and transfection supernatants were harvested 3-4 days post-transfection, aliquoted, and stored at −80°C. A/Victoria/210/2009 and A/California/07/2009 were further propagated in embryonated chicken eggs(44).

SARS-CoV-2 pseudotype viruses were generated on a vesicular stomatitis virus (VSV) pseudotype platform as previously described(45, 46). Briefly, VSV pseudotype virions bearing SARS-CoV-2 spike were produced by calcium phosphate transfection of 293T cells with 20 μg of pCG1 SARS-CoV-2 S Δ18 D614G expression plasmid encoding a codon-optimized SARS-CoV-2 spike gene with an 18 residue truncation in the cytoplasmic tail (provided by Stephen Pohlmann) (40) and a D614G substitution. 26 hours post-transfection, cells were infected with VSV-G pseudotyped VSVΔG-red fluorescent protein (RFP) at a MOI of ~1. 2-4 hours post infection, virus was removed, cells were washed twice with PBS, and media was replaced. 26-30 hours post infection, cell culture supernatant was harvested, clarified by centrifugation, and SARS-CoV-2 pseudotyped VSVΔG-RFP viruses were aliquoted and stored at −80°C. VSVΔG-RFP viruses bearing SARS-CoV-2 Beta (B.1.351) and Omicron (BA.1) spike were similarly produced.

### Recombinant proteins

Recombinant soluble trimeric HA (rHA) proteins were produced as previously described(47). Briefly, 293F cells were transfected with 1 ug/mL of expression plasmid encoding rHA constructs using 293Fectin in OptiMEM. Four days post-transfection, cell culture supernatant was clarified, and proteins were purified by gravity flow affinity chromatography with nickel-NTA agarose beads. HA proteins were buffer exchanged into PBS using 30K MWCO centrifugal filters. These HA proteins contained the following C-terminal modifications: a T4 Foldon trimerization domain, an AviTag for biotinylation, and a hexahistidine tag for purification. All C-terminal modifications were left intact for binding analyses. HA proteins used for influenza ELISAs were biotinylated using the site-specific biotin ligase, BirA, according to manufacturer guidelines (Avidity).

SARS-CoV-2 proteins were produced and purified in the same manner as the influenza HA proteins. A mammalian expression plasmid encoding the SARS-CoV-2 receptor binding domain (RBD) for the prototypic SARS-CoV-2/human/USA/USA-WA1/2020 strain was provided by Florian Krammer (Mt Sinai) (48). A similar expression plasmid encoding the RBD domain from a B.1.351 lineage SARS-CoV-2 virus was generated by restriction digest cloning, with a codon-optimized gBlock fragment (Integrated DNA Technologies) and a pCMV-Sport6 vector.

### Influenza HA enzyme-linked immunosorbent assays (ELISAs)

All ELISAs were performed on 96-well Immulon 4HBX flat-bottom microtiter plates coated with 0.5 ug of streptavidin per well, which was allowed to dry overnight at 37°C. Biotinylated recombinant HA diluted in 0.1% BSA in TBS + 0.05% Tween-20 was added to wells to serve as antigen. Plates were washed with PBS + 0.1% Tween-20 and then blocked with 1% BSA in TBS + 0.05% Tween-20 for one hour. Antibodies were serially diluted in 0.1% BSA in TBS + 0.05% Tween-20 and then added to wells for one hour. After washes with PBS + 0.1% Tween-20, HRP-conjugated mouse anti-human kappa (SB81a; Abcam) or lambda (JDC-12; Abcam) light chain secondary antibodies were diluted in 0.1% BSA in TBS + 0.05% Tween-20 and added to wells for one hour. After washes with PBS + 0.1% Tween-20, plates were developed by adding SureBlue TMB peroxidase substrate to wells for five minutes, after which the reaction was stopped with 250 mM hydrochloric acid. Plates were promptly read at an optical density of 450 nm on a microplate reader. Binding endpoint concentrations were defined as the of the lowest mAb dilution that gave an O.D. reading above ten times the plate background that day. Typical background O.D. values ranged from 0.04-0.06 across plates and over time, resulting in endpoint concentration cutoffs ranging from 0.4-0.6 O.D. units (450 nm).

### SARS-CoV-2 Spike enzyme-linked immunosorbent assays (ELISAs)

SARS-CoV-2 ELISAs were performed on 96-well Immulon 4HBX flat-bottom microtiter plates coated with 1 ug of antigen, which was incubated overnight at 4°C. On the day of the assay, plates were washed with PBS + 0.1% Tween-20 and then blocking buffer (3% goat serum and 0.5% milk in PBS + 0.1% Tween-20) was added to wells for one hour. Plates were again washed with PBS + 0.1% Tween-20, then antibodies were serially diluted 2-fold in blocking buffer and then added to wells for 2 hours. After washes with PBS + 0.1% Tween-20, HRP-conjugated mouse anti-human kappa (clone SB81a) or lambda (clone JDC-12) light chain secondary antibodies were diluted in blocking buffer and added to wells for one hour. After washes with PBS + 0.1% Tween-20, plates were developed by adding SureBlue TMB peroxidase substrate to wells for five minutes, after which the reaction was stopped with 250 mM hydrochloric acid. Plates were promptly read at an optical density of 450 nm on a microplate reader. Area under the curve was calculated in GraphPad Prism (v.9.2).

### Influenza virus neutralization assays

*In vitro* virus neutralization assays were completed as follows: 96-well flat bottom tissue culture plates were seeded with 2.5 x 10^4^ MDCK-Siat1-TMPRSS2 cells (49) per well a day before the assay in MEM supplemented with 10% heat-inactivated fetal bovine serum. On the day of the assay, ~1000 focus forming units (FFU) of virus were added to two-fold dilution series of antibody, and incubated at room temperature for one hour. Cells were washed twice with serum-free MEM to remove growth media, then 100 μL of virus/antibody mixture was added to wells. Plates were incubated at 37°C in 5% CO_2_ for 16-18 hours. Following incubation, media was aspirated from plates and cells were fixed with 4% paraformaldehyde (PFA) at 4C in the dark. PFA was then removed and 0.5% Triton X-100 was added to the wells for seven minutes. Triton X-100 was removed from wells and plates were blocked for one hour with 5% milk. Plates were then incubated with a mouse anti-influenza NP antibody (clone IC5-1B7) diluted in 5% milk for one hour, followed by incubation with peroxidase conjugated goat anti-mouse IgG diluted in 5% milk for one hour. Plates were then incubated with TrueBlue TMB substrate in the dark for one hour, after which substrate was removed and plates were allowed to dry before visualization and foci enumeration on an ImnmunoSpot S6 plate reader using the BioSpot program. Following the blocking, primary, and secondary steps, plates were hand washed 4 times with distilled water prior to the next step. Neutralization titers were expressed as 90% inhibitory concentration values, which were defined as the lowest concentration in the mAb dilution series that inhibited ≥ 90% of virus infectivity that was measured in virus only control wells.

### SARS-CoV-2 pseudotype virus neutralization assays

Neutralization assays using SARS-CoV-2 VSV pseudotype viruses were completed as previously described (45). Briefly, VeroE6/TMPRSS2 cells were seeded at 2.5×10^4^ cells/well in rat tail collagen coated 96-well plates. The following day, 300-500 FFU/well of VSVΔG-RFP SARS-CoV-2 pseudotype virus was mixed with serial 2-fold dilutions of antibody. A mouse anti-VSV Indiana G antibody, 1E9F9, was spiked into virus-antibody mixtures at 600 ng/mL. Virus-antibody mixtures were incubated for 1 hour at 37°C and then added to VeroE6/TMPRSS2 cells. 22 hours post infection, cells were washed with PBS, fixed with 4% paraformaldehyde, then blotted dry. Foci were visualized and counted on an ImnmunoSpot S6 plate reader using the FluoroSpot program. Half-maximal inhibitory concentration (IC_50_) was defined as the antibody concentration at which a 50% reduction in foci was measured relative to virus only control cells.

### Surface plasmon resonance

Binding kinetics of antibodies were measured by surface plasmon resonance (SPR) with a Biacore 3000 biosensor. Ni-NTA chips were loaded with 200 response units of his-tagged recombinant HA. The chip surface was regenerated at the beginning of each cycle with a 30 μL injection of 350 mM EDTA, followed by a 30 μL injection of 50 mM NaOH. Following regeneration, 5 uL of 0.5 mM NiCl2 was injected, followed by injection of His-tagged rHA at a flow rate of 5 μL/minute. Following a 1 min stabilization period, 200 μL of antibody was injected at a flow rate of 50 μL/minute for four minutes, followed by a dissociation phase of 20 minutes. Antibodies were injected at 800, 200, 50, 12.5, and 3.125 nM concentrations. Buffer-only injections were subtracted from antibody sensograms. All binding data were fit to simultaneous kinetics models provided in the BiaEvaluation software (v4.1). IgG were fit to 1:1 (Langmuir) binding models to determine an overall dissociation constant for the molecule (described in (50)).

## QUANTIFICATION AND STATISTICAL ANALYSIS

Specific statistical analyses are described in each figure legend. In all cases, data were graphed and statistical analyses were completed using GraphPad Prism (v.9.2). Data are represented as the mean of independent experiments, with error bars indicating the SEM. In some cases only the mean value is displayed, as indicated in the corresponding figure legend.

## Acknowledgements

This project has been funded in part with Federal funds from the National Institute of Allergy and Infectious Diseases, National Institutes of Health, Department of Health and Human Services, under Contract No. 75N93021C00015 and Grant No. 1R01AI108686. SEH holds an Investigators in the Pathogenesis of Infectious Disease Awards from the Burroughs Wellcome Fund. We thank J. Bloom (Fred Hutch) for providing MDCK-Siat1-TMPRSS2 cells. We thank S. Pohlmann for providing VeroE6/TMPRSS2 cells. We thank F. Krammer (Mt. Sinai) for providing the SARS-CoV-2 spike RBD expression plasmid.

## Author Contributions

Conceptualization, M.J.B. and S.E.H; experimentation, M.J.B., C.P.A, T.M.G., S.H.L., data analyses, M.J.B., SARS-CoV-2 neutralization assay development, P.B.; providing plasmids to express mAbs, P.W.; writing—original draft, M.J.B.; writing—review and editing, M.J.B, C.P.A, T. M.G, S.H.L, P.B., P.W., S.E.H.; supervision, S.E.H.

---

**Fig. S1.**
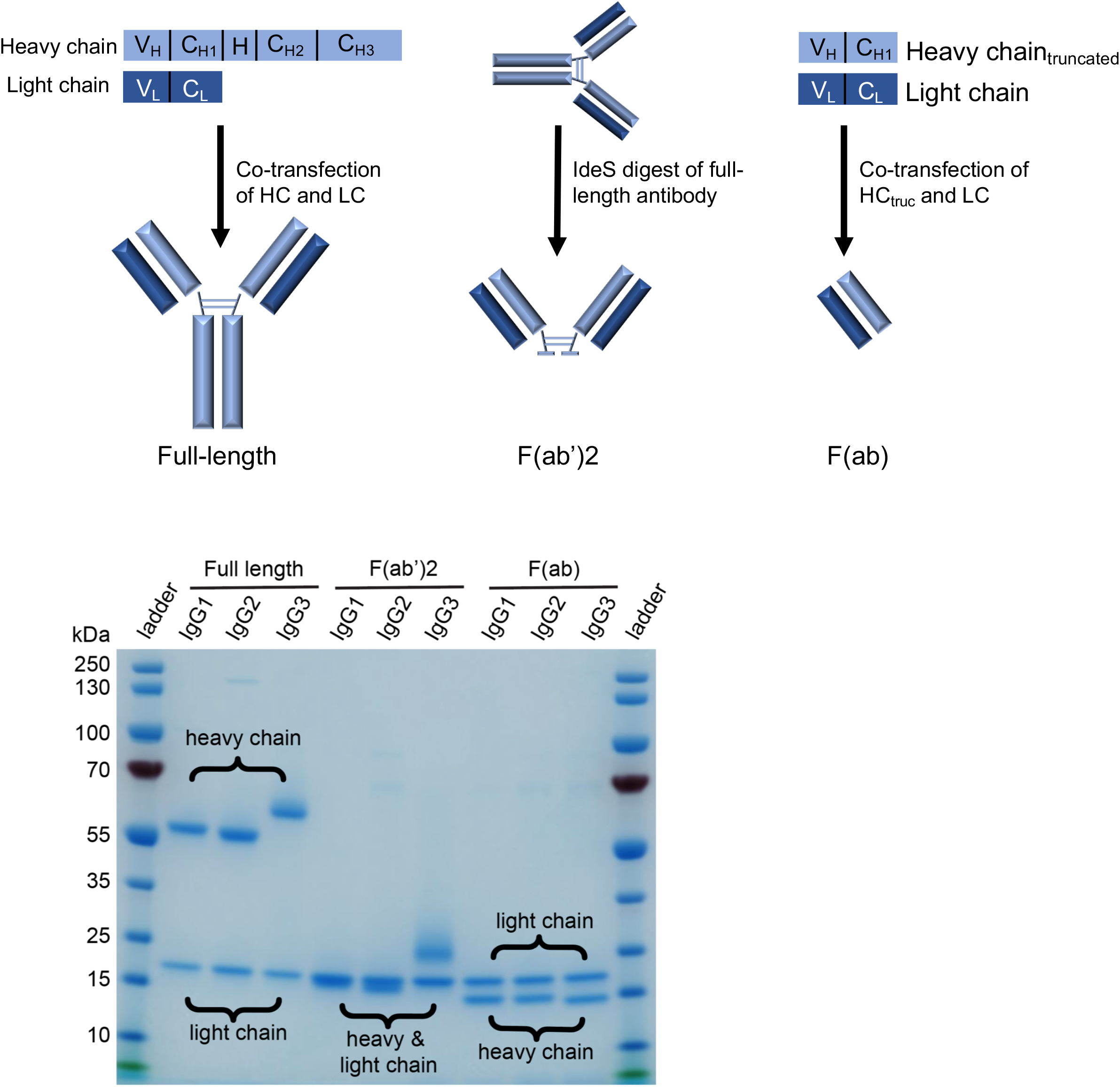
Antibody species generated in this study. (Top) Schematics depicting how full-length, F(ab’)_2_, and F(ab) monoclonal antibody species were generated in this study. (Bottom) Representative reducing SDS-PAGE analysis of one antibody, 10053-1G05, expressed as IgG1, IgG2, and IgG3 for each of the mAb species. Heavy and light chains are indicated.

**Fig. S2.**
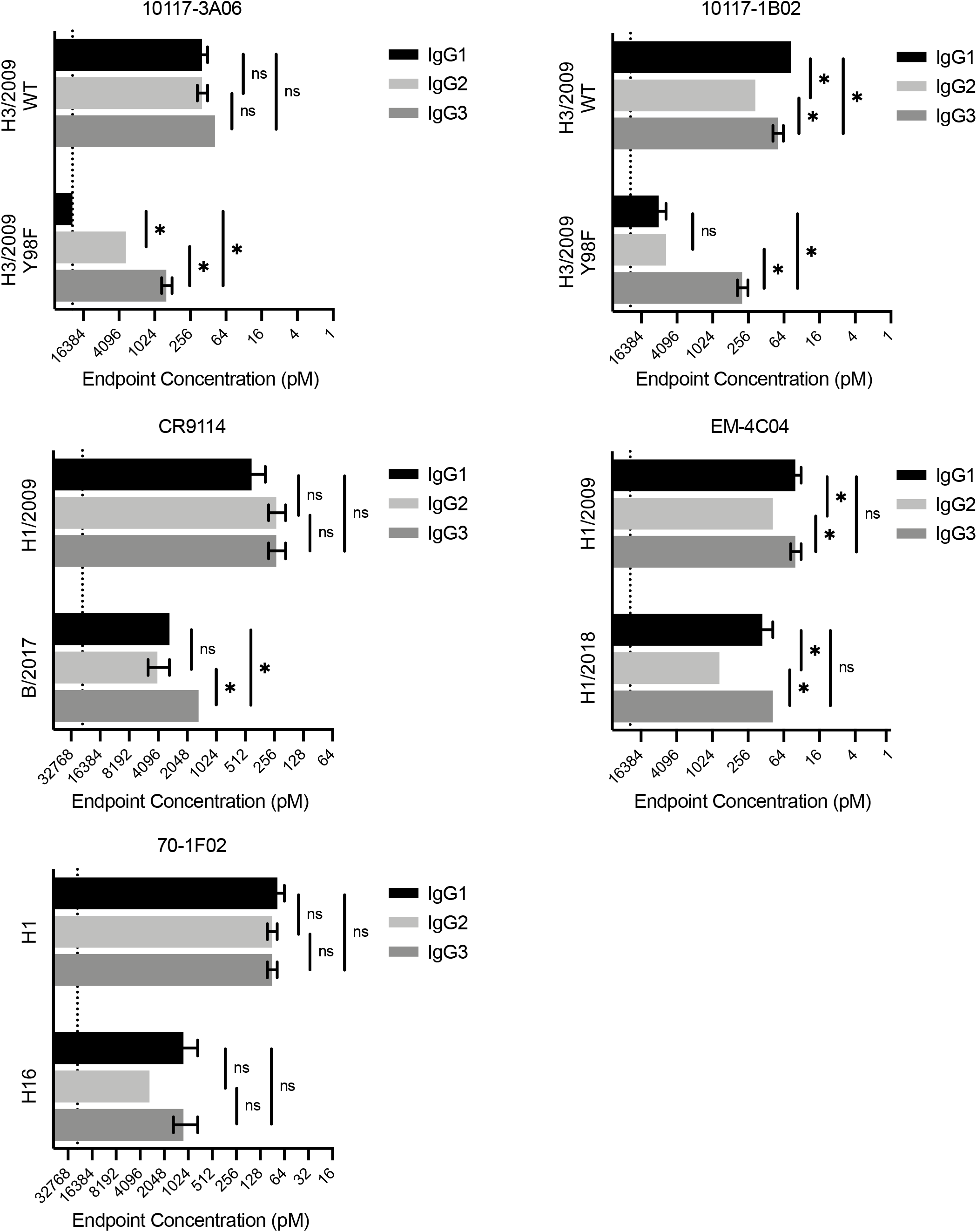
Influenza mAbs retain binding capacity to drifted influenza virus antigens when expressed as IgG3. ELISAs were completed with mAbs expressed as different IgG subclasses against antigens that reduce binding capacity of the mAb. ELISA binding values are shown as the mean endpoint concentration, with error bars indicating the SEM for 2-3 independent experiments. Dotted line indicates the limit of detection for the assay. Statistical comparison of the IgG subclasses for each antigen were completed using a one-way ANOVA on log2 transformed values with Tukey’s multiple comparisons post-test. *p<0.05

